# Natal and adult dispersal among four elephant seal colonies

**DOI:** 10.1101/2021.03.18.435977

**Authors:** Ramona Zeno, Richard Condit, Sarah G. Allen, Garrett Duncan

## Abstract

Dispersal plays a key role fostering recovery of endangered species because reoccupying a former range can only happen via dispersal. The northern elephant seal (*Mirounga angustirostris*) is a large, marine predator that was nearly exterminated in the 19th century by over-hunting. Once they were afforded protection from harvest, the species spread from a single remnant colony to reoccupy its former range. As colonies in central California were reestablished during the 1960s-1990s, tagged seals documented northward dispersal from southern California. The central California colonies are now large and well-established, and tagging programs at the four northernmost colonies allowed us to quantify the extent and direction of dispersal. Natal dispersal by females was highest from the southernmost colony at Piedras Blancas, where 61% of surviving females emigrated to breed. Dispersal from the other three colonies was much lower, 5.6% from SE Farallon Island, 10.3% from Año Nuevo, and 16.6% from Point Reyes. Adult dispersal of females, after breeding, was rare, with an annual rate < 2%. Juvenile dispersal is thus frequent in elephant seals, highest northward but also occurring southward, suggesting that continued expansion to new colonies throughout the west coast is probable.

## Introduction

Dispersal and immigration play an important role in the recovery of endangered species. The northern elephant seal was nearly exterminated during the 19th century by hunters collecting oil, but after hunting ended, it recolonized much of its historic range [[1–3]]. In previous work during the 1970s, high numbers of immigrant females at the Año Nuevo colony in central California were documented, and population growth was fueled by immigration [[4–7]]. The flow of dispersal among colonies was northward, from large breeding populations in southern California toward Año Nuevo [[8, 9]]. Since the 1970s, additional colonies have formed in central California, again due to steady dispersal from the south [[3, 7]].

The elephant seal’s rapid recovery contrasts with other pinnipeds such as the Guadalupe fur seal (*Arctocephalus townsendi*) and New Zealand sea lion (*Phocarctos hookeri*). Both species were hunted to scarcity a century ago, and their populations remain restricted. Strong site fidelity and poor dispersal have hampered recovery [[10, 11]]. Quantifying dispersal thus may provide a tool to predict whether populations can recover.

Now that the elephant seal colonies in central California colonies have reached large size and are monitored regularly [[3, 12, 13]], we can revisit the importance of immigration, this time with the intent of quantifying the rate in all directions. To quantify dispersal and immigration rates, we tagged pups at four different colonies during 1998-2000 and maintained consistent sighting efforts at all sites through 2008. This is the first study to quantify rates of immigration among several elephant seal colonies.

## Materials and Methods

### Study sites and colonies

The four colonies are 35.7° N to 38.9° N latitude on the California coast and are the northernmost large colonies of the species (Fig 1). At all four sites, female elephant seals gather in large groups on flat sand beaches and give birth to a single pup per year from December-February. Pups are weaned an average of 25 days after birth when mothers depart to forage, and weanlings are easily approached and tagged on the beach before they go to sea [[14]].

**Figure 1.**
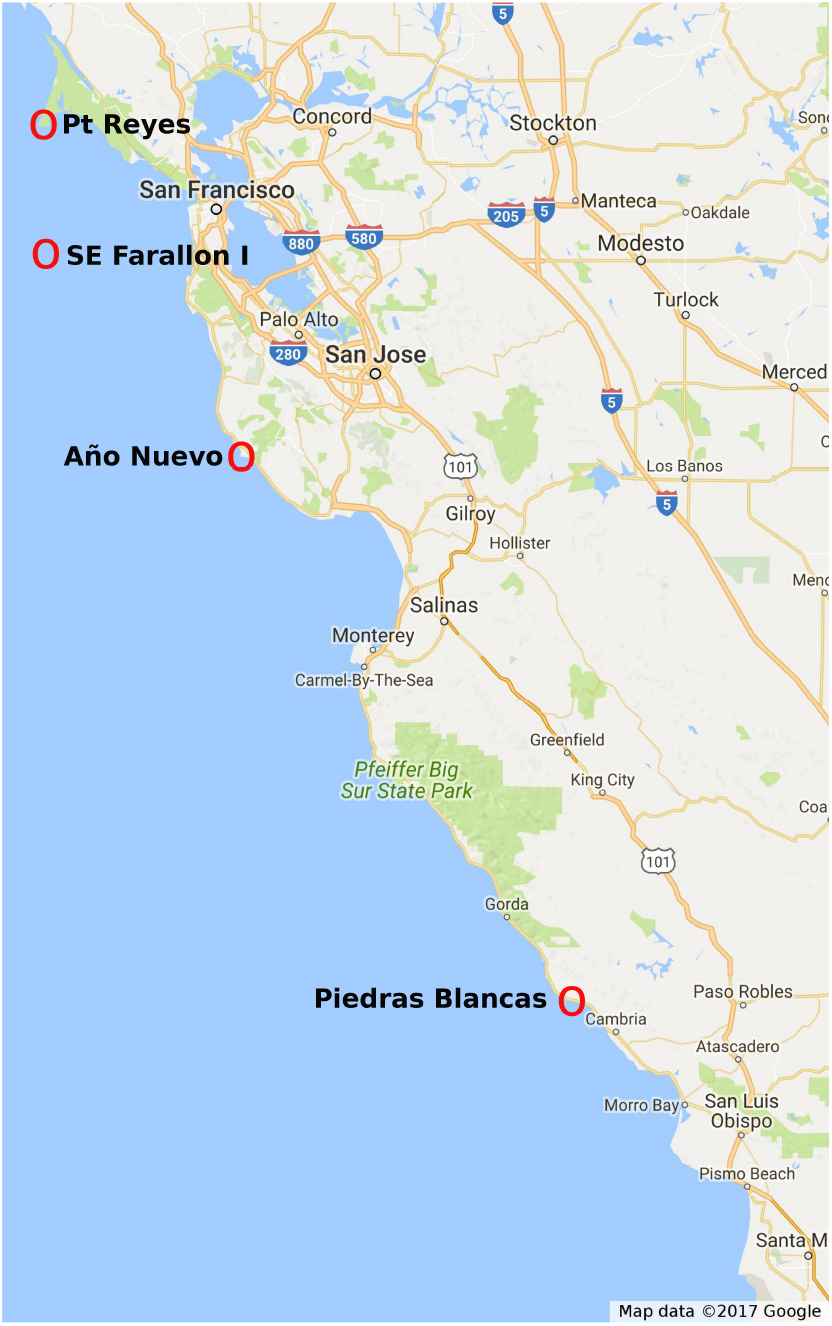
Location of the four elephant colonies (Table 1).

### Tagging and merging data

Plastic cattle tags were inserted in the interdigital webbing of the hind flippers, usually one on the left flipper and one on the right, and tag type and tagging method were identical at all four sites [[12, 15, 16]]. Tags were color-coded by colony and bore unique numbers. Although seals have been tagged for many years, we focus on three cohorts: females tagged as pups in 1998, 1999, and 2000, a period when all four colonies had large numbers tagged and followed by many years of consistent resighting effort.

The resight effort was concentrated in the winter breeding season, when females with their pups hold their ground and allow observers close. Tag numbers were read with binoculars by approaching within 3-5 m of seals or with telescopes from 5-20 m, the latter from dunes, cliffs, or platforms above the seals. We carried out thorough searches for tagged females at each colony and then collected all sightings of the three focal cohorts from 2001-2008. Since females first breed from age 3 to 5 years old, this sample provided a consistent opportunity to observe females from all cohorts at all four colonies. We assumed any female age 3 or older observed during the breeding season was breeding because 97.5% of adult females in the colony give birth [[6]].

Double-tagged females, carrying two different numbers, required considerable effort in cross-referencing sighting data to make sure each record of a tag was attributed to the correct female. Prior to this study, sightings of doubly-tagged seals at foreign colonies (other than the tagging location) were sometimes attributed to two different animals, which happens when the two tag numbers are not seen during the same sighting. Once tags were correctly assigned to each individual, we constructed a matrix of all seal sightings in all breeding years. In these cohorts, we never saw a female at two different locations during the same year, thus a single record per year was sufficient to indicate each animal’s location.

### Natal dispersal

Both juvenile and adult elephant seals are site tenacious [[9]], however, young elephant seals colonize new colonies more often than adults [[5, 17, 18]]. We thus considered juvenile and adult dispersal separately. To estimate natal dispersal, define *T_i_* as the total number of females tagged at colony *i*, and *B_i_* the number of those observed breeding at any time in the future. *B_i_* can be separated into four groups to indicate where each female was first observed breeding: *b_ij_* is the number born at colony *i* and first observed breeding at colony *j*. The ratio 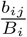 is a measure of natal dispersal rate from colony *i* to colony *j*. Many of the original *T_i_* females tagged were never seen breeding, but those do not enter into the calculation, since we know nothing about their fate; they may have dispersed then died before a breeding observation, or simply died before dispersing. All calculations were done summing across the three cohorts to provide a more robust sample size (Table 1). Credible intervals on the number migrating were calculated using the beta distribution to describe the posterior sampling distribution of the number migrating, *b_ij_*, out of the total, *B_i_* [[19]]. This assumes *b_ij_* is binomially distributed, though with four different colonies it is actually a multinomial, nevertheless, the beta limits based on the binomial are good approximations [[20]].

**Table 1.** Sample size of females from 1998-2000 cohorts. Tagged gives the number of female pups to which permanent plastic tags were deployed. Breeding includes the total number of those tagged that were observed breeding at any of the four colonies. Dispersing are those first observed breeding at a colony different from their birth location.

| Colony | Tagged $T_i$ | Breeding $B_i$ | Dispersing $\sum_{i \neq j} b_{ij}$ |
| --- | --- | --- | --- |
| Pt Reyes | 216 | 30 | 5 |
| SE Farallon | 154 | 18 | 1 |
| Año Nuevo | 616 | 107 | 11 |
| Pd Blancas | 448 | 101 | 62 |
| Total | 1434 | 256 | 79 |

### Adult dispersal

Once a female breeds at one colony, she might then move to another in a subsequent breeding attempt. To evaluate adult dispersal, we first documented all females observed breeding at two different locations. This turned out to be such a small number that we present each animal’s full breeding history. As an estimate of adult dispersal, we then consider a restricted sample: those females observed breeding in consecutive years. We counted all cases where a female dispersed between those observations, and all cases where she did not. We could tally this for every pair of colonies *i* to *j*, but there were so few we present them as a single fraction dispersing. In addition, we present the percentage of all breeding females that were observed dispersing as adults as an estimate of the lifetime adult dispersal rate.

### Juvenile-adult dispersal correlation

The sample of individually-marked females and their lifetime movements allow a test for correlation between juvenile and adult dispersal. That is, are animals which dispersed as juveniles more likely or less likely to move as adults? The sample of 226 females observed breeding twice was divided into two groups, those dispersing as juveniles and those not. For each group, the fraction that dispersed again as adults was calculated.

## Results

### Natal dispersal

Of all females tagged as pups during 1998-2000 at the four colonies, 256 were later seen breeding, and 79 of those (31%) dispersed as juveniles to start breeding at a foreign colony. The proportion emigrating, however, varied greatly from colony to colony. At the three northerly colonies, 83-94% of females were site tenacious and returned to breed where they were born (Table 2). Those three rates were statistically indistinguishable, and the combined mean was 89%. In contrast, only 39% were site tenacious at the southern colony, Piedras Blancas. More than half the Piedras-Blancas females emigrated, a rate five times higher than the other sites, and 95% credible intervals were widely separated (Table 2). For three sites, most emigrants went to Año Nuevo; from Año Nuevo, most went to Point Reyes (Table 2).

**Table 2.** Juvenile dispersal rates. Bold-face entries are percentages of animals born at one site (the row) and first observed breeding at a second site (the column). Only animals observed breeding at least once are included in the percentage (samples are in Table 1); entries across a row add to 100%. Below each bold-face entry are 95% credible intervals in parentheses. The diagonal entries are non-dispersal, where birth and breeding colonies are the same.

| Birth colony | Breeding colony |  |  |  |
| --- | --- | --- | --- | --- |
|  | Pt Reyes | SE Farallon | Año Nuevo | Pd Blancas |
| Pt Reyes | **83.3**<br>(66.3,92.5) | **0.0**<br>(0.0,11.2) | **13.3**<br>(5.5,29.8) | **3.3**<br>(0.8,16.7) |
| SE Farallon | **0.0**<br>(0.0,17.6) | **94.4**<br>(74.0,98.7) | **5.6**<br>(1.3,26.0) | **0.0**<br>(0.0,17.6) |
| Año Nuevo | **6.5**<br>(3.3,12.9) | **2.8**<br>(1.0,7.9) | **89.7**<br>(82.5,94.1) | **0.9**<br>(0.2,5.1) |
| Pd Blancas | **12.9**<br>(7.7,20.8) | **15.8**<br>(10.0,24.2) | **32.7**<br>(24.3,42.3) | **38.6**<br>(29.7,48.4) |

### Adult dispersal

Dispersal after breeding began was rare. We observed 270 chances to disperse over consecutive years, and five dispersal events, for a rate of 1.9% y^−1^ (95% credible interval 0.8-4.2%). Those five events included just four females, because female GO894 dispersed twice (Table 3). The four females observed moving over consecutive years included two born at Año Nuevo and two at Piedras Blancas, but their movements included all four colonies (Table 3). Expanding the sample to include all 156 females seen breeding more than once, not necessarily in consecutive years, there were seven additional females who moved as adults (Table 3). Thus, over a lifetime, we observed 11 of 156 adult females dispersing, suggesting a lifetime rate of 7.1%.

**Table 3.** Adult dispersal by elephant seal females from 1998-2000 tagging cohorts, showing lifetime records of all 11 seen breeding at two or more colonies. The years 1998-2000 show where each was born, and 2002-2007 where they bred every time they were observed; AN=Año Nuevo, PR=Pt Reyes, PB=Piedras Blancas, SEF=SE Farallon Island. The first four animals included the only cases where dispersal was confirmed by sightings in consecutive years at two different colonies; those were used in estimating annual dispersal probability. The remaining seven dispersed but had a sighting gap between. Eight of the 11 also dispersed as juveniles, between birth and first breeding. The other three were not juvenile dispersers as tallied in Table 2, because they were primiparous at their natal colonies.

| Female | Birth colony |  |  | Breeding colony |  |  |  |  |  |
| --- | --- | --- | --- | --- | --- | --- | --- | --- | --- |
|  | 1998 | 1999 | 2000 | 2002 | 2003 | 2004 | 2005 | 2006 | 2007 |
| GO894 |  | AN |  | AN | AN | AN | PB | AN |  |
| GO973 |  | AN |  |  | PR | AN | AN | AN |  |
| W1114 | PB |  |  | SEF | SEF | AN |  |  |  |
| W1198 |  | PB |  | AN | PB |  |  |  |  |
| GO123 | AN |  |  | AN | AN |  | PB | PB |  |
| GO229 |  | AN |  |  | AN |  |  | PR |  |
| W1102 | PB |  |  |  | AN |  | PB | PB |  |
| W1196 |  | PB |  | AN |  |  |  |  | PB |
| W1310 |  | PB |  |  | PR |  | PB |  |  |
| WX 50 |  |  | PB |  | AN |  |  | PR | PR |
| WX251 |  |  | PB |  |  | AN |  |  | PR |

### Juvenile-adult comparison

Females who dispersed as juveniles were six-times more likely to later disperse as adults (8 of 46 juvenile dispersers became adult dispersers, compared to 3 of 110 non-dispersing juveniles, including only females seen breeding at least twice). Concomitantly, most of the adult dispersers (8 of 11) had already dispersed as juveniles. Of the eight dispersing both as juveniles and adults, five moved back to where they were born, but the other three moved to yet another colony, i.e. were born at one colony then bred at two other colonies (Table 3).

## Discussion

Natal dispersal was frequent in elephant seals, reaching 61% from one colony, while breeding dispersal was rare, averaging just 2% per year. This is a common pattern in vertebrates, for example, Paradis et al. [21] found that 61 of 69 British bird species disperse further as juveniles than as adults, and seabirds and other pinnipeds are similarly more site tenacious after breeding begins [[22–25]].

Juvenile dispersers were more likely to disperse again as adults. This might suggest a genetic proclivity toward dispersal, or perhaps it means that a female who discovers a second colony subsequently has two sites from which to choose. Interestingly, though, three of the 11 adult dispersers moved not back to their birthplace, but to a third colony. Finally, despite elevated likelihood of dispersal as adults, those juvenile dispersers were site tenacious. Once choosing a new colony, > 80% stayed there throughout their lives [[26]].

We quantified earlier observations of a northward bias in natal dispersal by elephant seals [[8]], finding that 61% of females emigrated northward from Piedras Blancas, while only 1% emigrated southward from Año Nuevo and 17% from Point Reyes. A directional bias is not considered in broad reviews of natal dispersal [[21, 27]], but it is of paramount importance in elephant seals. The remnant left behind by hunters in 1890 was at the south end of the current range, on Guadalupe Island in Mexico. This appears simple good fortune, when coupled with the proclivity toward northward natal dispersal. Had the remnant colony been at the north end of the range, the species may not have rebounded so rapidly.

During the 1970s when northward dispersal was first documented [[8]], the most likely hypothesis for it was simple mass action. Colonies further south expanded sooner, and during the 1960s and 1970s, when they were much larger than northern colonies, more animals would be moving northward than southward even without a directional bias. Such a mass-action effect would weaken, though, as the northern colonies grew. Our current observations, however, suggest that individual animals move northward with higher probability than southward. Moreover, since the 1970s, a new source of understanding has emerged from detailed studies of feeding migrations. Elephant seals travel northward or northwestward great distances to feeding grounds [[28–31]], and this means that juveniles from southern California migrate past, and sometimes not far from, colonies in central California. Observations of juveniles from southern colonies at Año Nuevo during the annual molt are thus unsurprising, and these prospecting juveniles sometimes breed at Año Nuevo [[32]].

There is, however, an important caveat about directional dispersal. Females are not always observed when present on a breeding colongy. At Año Nuevo, we estimated detection probability at 70% per year for breeding age females [[33]]. If this detection probability varied across colonies, then estimates of dispersal would be biased. From the data currently available, we cannot test for variation in detection probability, so it is possible that the difference between northward and southward dispersal rates is an artefact of detection.

Regardless of the directional bias, natal dispersal was common, averaging 11% at three colonies and 61% from Piedras Blancas. Since the elephant seal population continues to expand [[3]], it remains likely that new colonies will form anywhere on remote beaches of the west coast. We recommend that management of coastal systems include the possibility of new colonies forming, such as the small ones known from northern California to British Columbia [[34, 35]]. New colonies may encroach upon breeding habitat of imperiled species such as the western snowy plover (*Charadrius alexandrinus nivosus*), which breeds on a few undisturbed California beaches. In Point Reyes, elephant seals and plovers compete for limited undisturbed space [[36]]. Rare plants native to coastal dunes are also potentially at risk of elephant seals seeking unoccupied haul out sites to molt or breed. Moreover, dispersal patterns affect dynamics of the existing colonies, some of which continue to grow while others are declining [[3, 13, 37, 38]]. Further monitoring of dispersal is thus important for anticipating future population changes in the species.

## Acknowledgments

We are grateful to B. Hatfield USGS Western Ecological Research Center for cohort sizes and many female sightings at Point Piedras Blancas, and to B.J. Le Boeuf and D.P. Costa for organizing and supervising the tagging program at Año Nuevo for many years. We also thank many colleagues, students, and volunteers for years of tagging and tag sighting. H. Huber, P. Pyle, and R. Bradley provided long-term support at the Farallon Islands. D. Adams, D. Roberts; D. Press at Point Reyes; and K. Karako Piedras Blancas. P.A. Morris was crucial in maintaining consistent tag sighting effort at Año Nuevo. The research was supported by the Tagging of Pacific Pelagics program with grants from the National Ocean Partnership Program (N00014-02-1-1012), the Office of Naval Research (N00014-00-1-0880, N00014-03-1-0651 and N00014-08-1-1195), International Association of Oil and Gas Producers contract JIP2207-23, the Moore, Packard, and Sloan Foundations. The surveys conducted on the Farallon Islands and Point Reyes were authorized by National Marine Fisheries Service permit #373-1575, and at Año Nuevo by #87-1743-04. Work was conducted under Marine Mammal Research Permits 347, 404, 684, 704, 774-1437, 774-1714, and 14097; and National Marine Sanctuary Permits GFNMS/MBNMS/ CINMS-04-98, MULTI-2002-003, MULTI-2003-003, and MULTI-2008-003. All authors were involved in the collection of data and approved the final manuscript.

